# A Graph Neural Network Approach to Investigate Brain Critical States Over Neurodevelopment

**DOI:** 10.1101/2024.10.09.617395

**Authors:** Rodrigo M. Cabral-Carvalho, Walter H. L. Pinaya, João R. Sato

## Abstract

Recent studies show that functional resting-state dynamics may be modelled by lattice models near criticality, such as the 2D Ising model. The Ising temperature, which is the control parameter dictating the phase transitions of the model, can provide insight into the large-scale dynamics and is being used to better understand different brain states and neurodevelopment. This period is categorized by intricated changes in the microcircuits to consolidate networks. These changes influence the macroscopic brain dynamics and also its functional relations, which can be observed in functional Magnetic Resonance Imaging (fMRI). Therefore, this work investigates neurodevelopment through a novel method to estimate the Ising temperature of the brain from fMRI data using functional connectivity and Graph Neural Networks (GNNs) trained on Ising model networks. The main finding indicates a statistically significant negative correlation between age and temperature for typically developing children (𝑟 =− 0. 48, 𝑝 < 0. 0001) and also children with Attention deficit/hyperactivity disorder (ADHD) (𝑟 =− 0. 49, 𝑝 < 0. 0001). This study suggests that the brain gets distant from criticality as age increases, leading to a more ordered state.

**Author Summary:** In this study, we employed Graph Neural Networks to investigate functional connectivity patterns of resting-state fMRI data using the 2D Ising model as a theoretical framework to access critical dynamics. Our key finding is the significant decrease in Ising temperature with age in both typically developing children and those with ADHD, indicating a developmental transition from a high-entropy, disordered state to a more ordered, decreasing the availability of dynamical states that the system can occupy and transit in different conditions. Moreover, this decrease in temperature is crucial as it implies that the brain is moving away from criticality, leading the system to a more sub-critical regime. The correlation coefficients across the groups showed no significant differences, indicating the same developmental trajectory.

## 1 Introduction

The brain is structured as a vast, astronomical-scale network, governed by complex non-linear biochemical interactions and dynamically influenced by environmental information. This complex organization gives rise to various microscopic and macroscopic spatiotemporal patterns and cognitive, emotional, motor, and sensorial behaviours (Kelso, S., 1995). The remarkable complexity of brain dynamics has been attracting the interest of physicists eager to model a set of equations that can give some perspective on self-organization and dynamical patterns. Thus, marvellous contributions propose brain dynamics as a collective process from microscopic components using tools from statistical physics and complex systems (Werner, 2007). One of the most prominent results shows evidence that resting state brain networks extracted from functional Magnetic Resonance Imaging (fMRI) BOLD-signal are statistically indistinguishable from ones extracted from the 2D Ising model (Fraiman et al., 2009), a mathematical and computational model from statistical physics to study phase transitions, showing that the macroscopic brain dynamics functionally operates near second-order phase transition elucidated by the 2D Ising model (Chialvo, 2010). This model explains magnetization as an emergent phenomena, known as spontaneous magnetization, using only collective microscopic interactions, which are controlled by a control parameter (Ising temperature). These emergent patterns appear at a specific interval of values of the control parameter, which is the critical point where the system transitions between two or multiple states (Beggs, 2008; Haimovici et al., 2013). Consequently, the concept that the brain operates near a critical point introduces the term "Critical Brain."

Despite the significant differences between the 2D Ising models, which consist of a simple two-dimensional lattice with local direct interactions ruled by a Hamiltonian, and the complex dynamics of the brain, it can provide some insights about whole-brain dynamics from the statistical physics perspective. It’s important to note that many studies have focused on developing improved Ising models for brain dynamics, for example, by incorporating structural data (Nuzzi et al., 2020). These models use techniques such as the pairwise maximum entropy model (PMEM) and maximum likelihood estimation to infer parameters, such as the interaction matrix J and the external magnetic field h, from the data (Ezaki et al., 2017). However, using PMEM when the number of ROIs is large remains an issue (Ezaki et al., 2020). Therefore, this present work focuses on the simpler 2D Ising model, for the sake of creating an approach based on machine learning that operates at the connectivity level for inferring the temperature for 333 ROIs (i.e., a large number of ROIs) with a light computational cost for scalability over large datasets. Deep learning models can incorporate various patterns and generalize over different dynamical systems (Legaard et al., 2023); thus, this work proposes an approach that can be extended to a deep learning model that learns from a variety of biologically informed and improved Ising models.

A geometric deep learning approach was employed to extract the complex features of connectivity information related to the Ising temperature in the network structure. This specialized Machine Learning algorithm approach operates on non-Euclidean spaces like networks. One of the most common approaches is the Graph Neural Network (GNN), which was trained using simulated 2D Ising model networks to predict the control parameter (i.e., Ising temperature) that originated the dynamics and, consequentially, the connectivity graph. Therefore, once this geometric deep learning method was trained to correctly predict the Ising temperature, it was applied to estimate the temperature across neurodevelopment (ages 8 - 22 years) from rs-fcMRI graphs.

This period is heavily categorized by intricate changes in the microcircuits across the brain, especially in the cerebral cortex, to consolidate and create more cohesive networks (Paolicelli et al., 2011; Zuo et al., 2017). These significant changes influence the macroscopic brain dynamics and also its functional relations, which was observed in rs-fcMRI data, demonstrating mechanistic alterations on brain dynamics at the network level, representing a wide variety of neural patterns enabled by more flexible dynamics (Menon, 2013; Sydnor et al., 2021) and producing effects on the functional networks topological properties (Sato et al., 2016; Ernst et al., 2015; Betzel et al., 2014; Grayson & Fair, 2017; Soman et al., 2023). Thus, the absence of topological characterization of networks limits the analysis of studies where structural connectivity experiences significant changes. Therefore, due to the limitations of the 2D Ising model with constant interactions and first neighbors coupling, the difference in critical state transitions using functional connectivity networks can only be assessed using Ising temperature.

In this context, this work examines neurodevelopment using Ising temperature to address how critical states relate to the formation and evolution of functional networks during neurodevelopment by the change in the control parameter. Indicating a departure from criticality over age, moving from a disordered phase to a more ordered one, where neural activity becomes less random and more predictable. Even though limited by a lack of topological characterization, the present framework inaugurates a possibility of temperature estimation with a computational light approach for large datasets, with many ROIs (i.e., 333), and extendable to diverse training sets extracted from various dynamical models to infer control parameters.

## 2 Results

### 2.1 Estimating Ising temperature during neurodevelopment

Once the GNN was trained on the Ising simulations over 200 time points, the Ising temperature of the rs-fMRI was estimated for Typically Developing Children and ADHD symptoms. Thus, the difference between the estimated Ising temperature T and the critical temperature *Tc* (T - Tc) was calculated. Hence, Pearson’s and Spearman’s Correlation were calculated, evidencing that both groups show a significant negative linear relationship between estimated temperature and age. For Typically Developing Children (Pearson = -0.48, *p < 0.0001*; Spearman = -0.47, *p < 0.0001*). For ADHD symptoms (Pearson = -0.49, *p < 0.0001*; Spearman = -0.47, *p < 0.0001*).

The negative trend for the GNN trained on the 2D Ising model for 2000 time points was consistent with this result, which showed statistical significance and can be found in the supplementary materials (Table A). Moreover, both groups are below the critical temperature *Tc* with a linear decrease in Ising temperature over neurodevelopment (Fig. 1). Additionally, potential relationships between Ising temperature and key graph topology properties were analyzed using Pearson’s correlation, with the results presented in the Supplementary Material (Table B).

**Figure 1.**
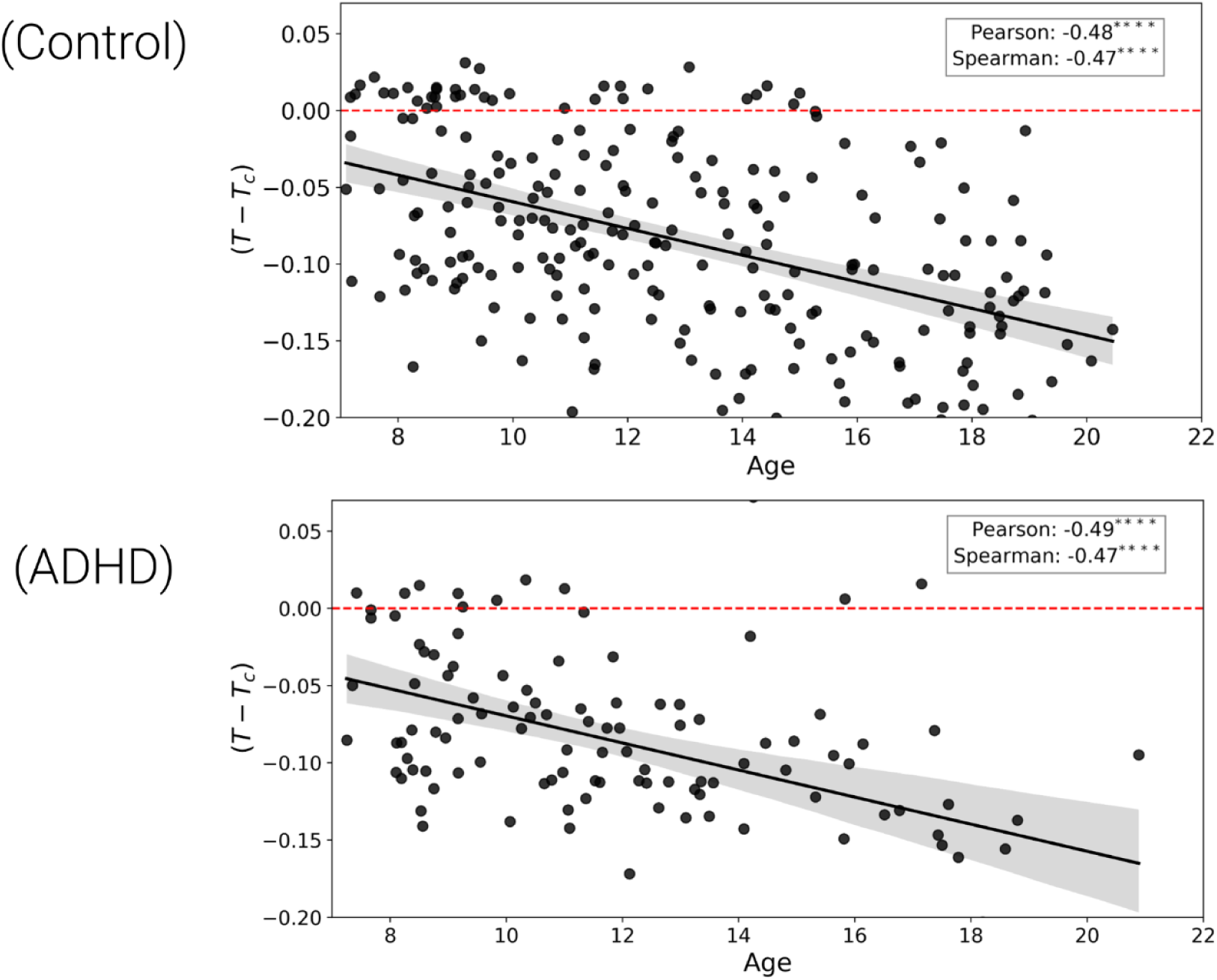
Estimated Ising temperature T minus critical temperature *Tc* plotted against age. The top plot (Control) shows typically developing children, and the bottom plot (ADHD) shows children diagnosed with Attention Deficit Hyperactivity Disorder (ADHD). The solid black line in each plot represents the linear trend between T - Tc and age, with a shaded region indicating the standard deviation. In both plots, a significant negative correlation between T - Tc and age is observed, as indicated by Pearson’s correlation and Spearman’s rank correlation values with corresponding asterisks denoting the level of statistical significance (****𝑝 ≤ 0. 0001). The dashed red line represents the critical temperature Tc as a reference.

The age was divided into seven groups of 2 years (7-9, 9-11, 11-13, 13-15, 15-17, 17-19, 19-21), and the distance of the estimated Ising temperature T to critical temperature *Tc* |T - Tc| was calculated for each age group. As can be observed in Fig. 2, both typically developing children and subjects with ADHD symptoms are getting distant from criticality over neurodevelopment. Even though the median distance from the critical point always increases, not all groups are statistically different. Therefore, as expected, age groups distant in the age span show statistical significance. A two-sample t-test was used to assess differences in the distance to the critical temperature between age groups; the p-value for pairwise comparison between each group can be seen in Table 1.

**Figure 2.**
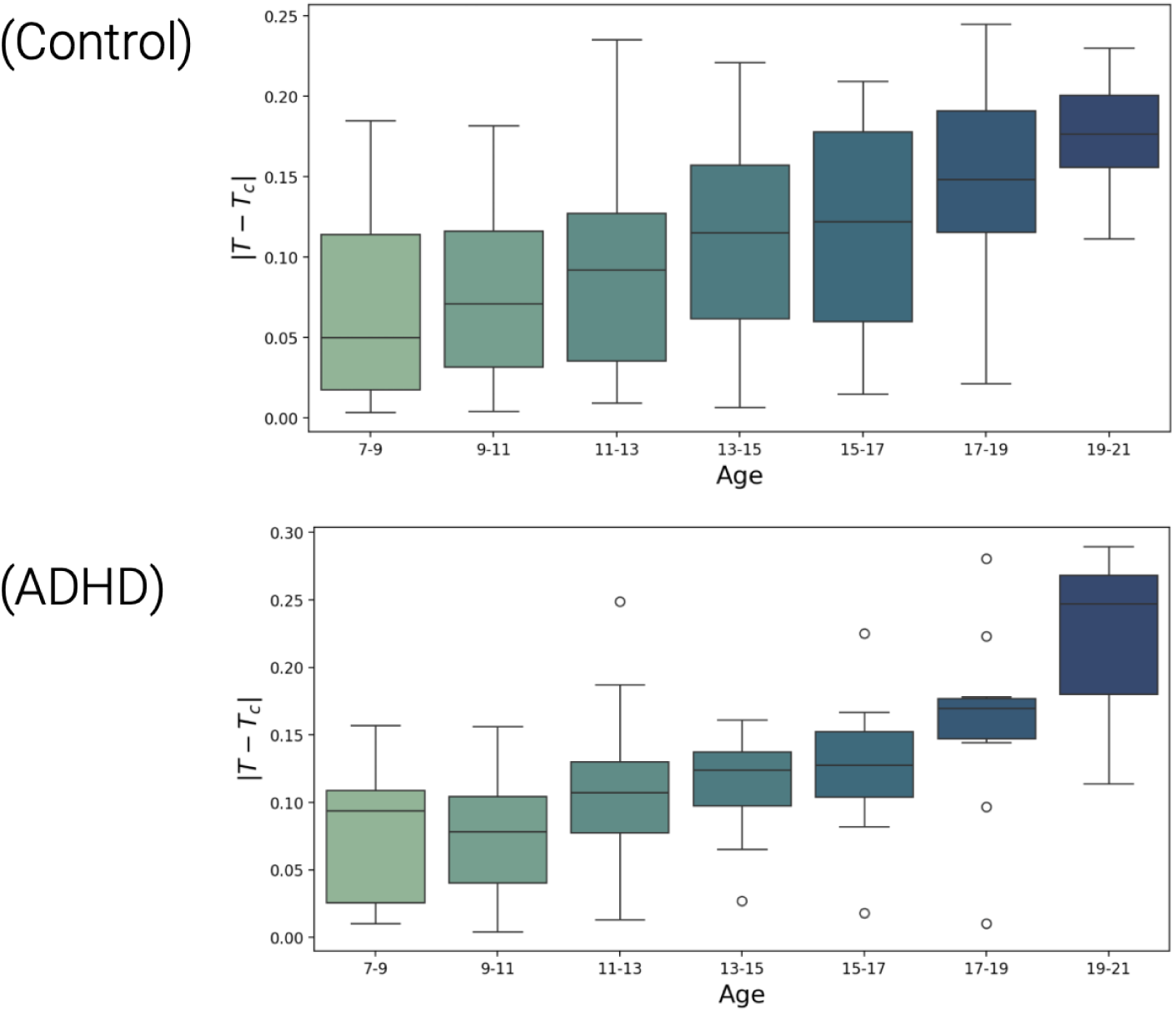
The box plots represent the distance from the estimated Ising temperature T to the critical temperature Tc (|T - Tc|) across age groups for both typically developing children (Control) and children with ADHD. The boxes show the interquartile range (IQR) with medians, and the error bars indicate data within 1.5 times the IQR. Circles represent outliers, which fall outside this range and are particularly noticeable in the ADHD group for older age groups.

**Table 1.**
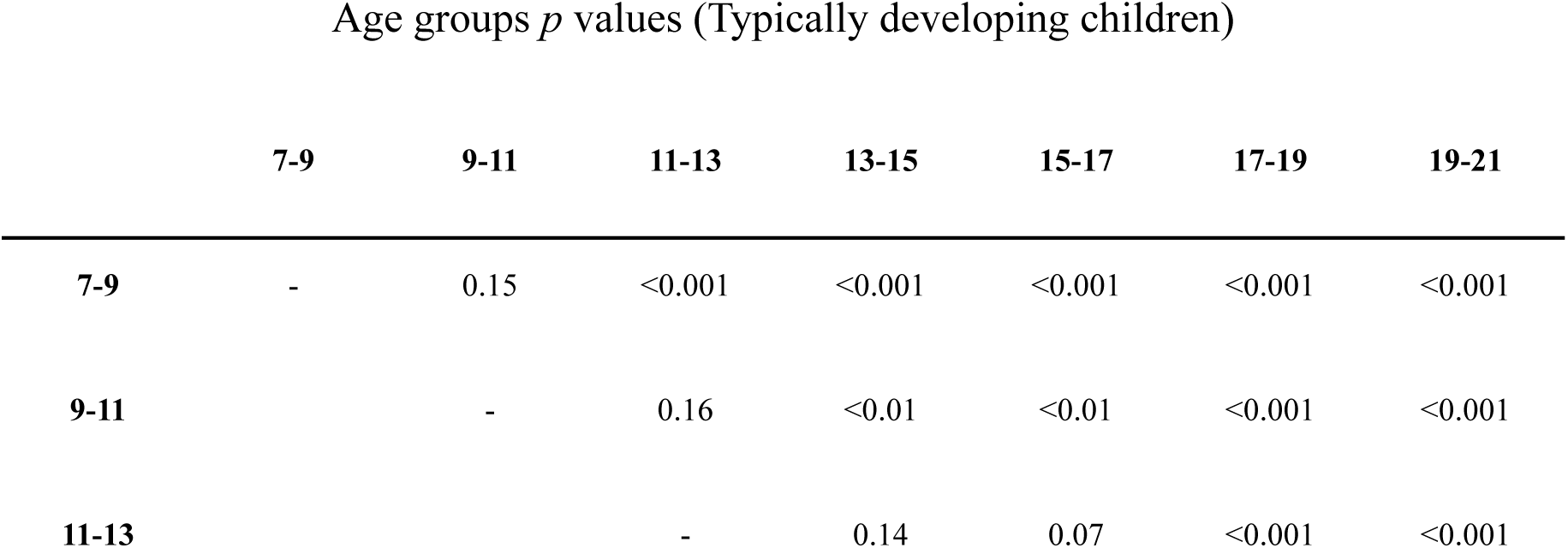
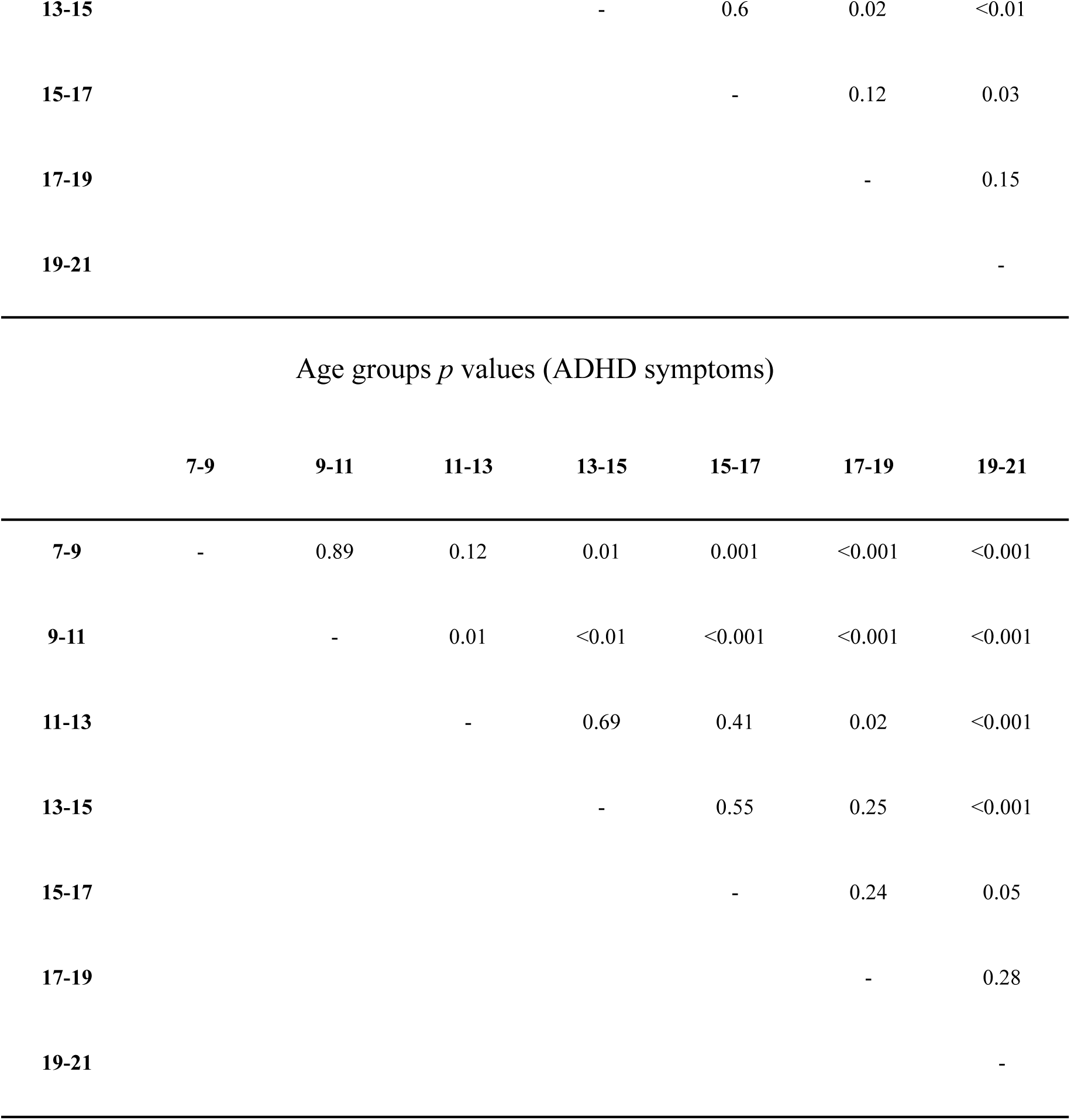
The paired t-test p-values results for the distribution of the distance from the estimated Ising temperature from the critical temperature for pairwise age groups. Top table - p-values for Typically developing children. Bottom table - p-values for ADHD symptoms.

### 2.2 Group differences

A bootstrap analysis was conducted to compare the distributions of the correlation coefficients for each group to evaluate the difference between the groups’ correlation coefficients. This statistical technique involves repeatedly resampling the data with replacement to create a distribution of correlation coefficients, allowing for a more robust comparison between the groups. The analysis revealed that the presence of ADHD symptoms does not significantly affect the observed decrease in Ising temperature with age (Pearson’s Correlation, *p = 0.5*; Spearman’s Correlation, *p = 0.6*), indicating that the trend of the brain moving away from criticality as it matures is consistent regardless of ADHD status (Fig. 3).

**Figure 3.**
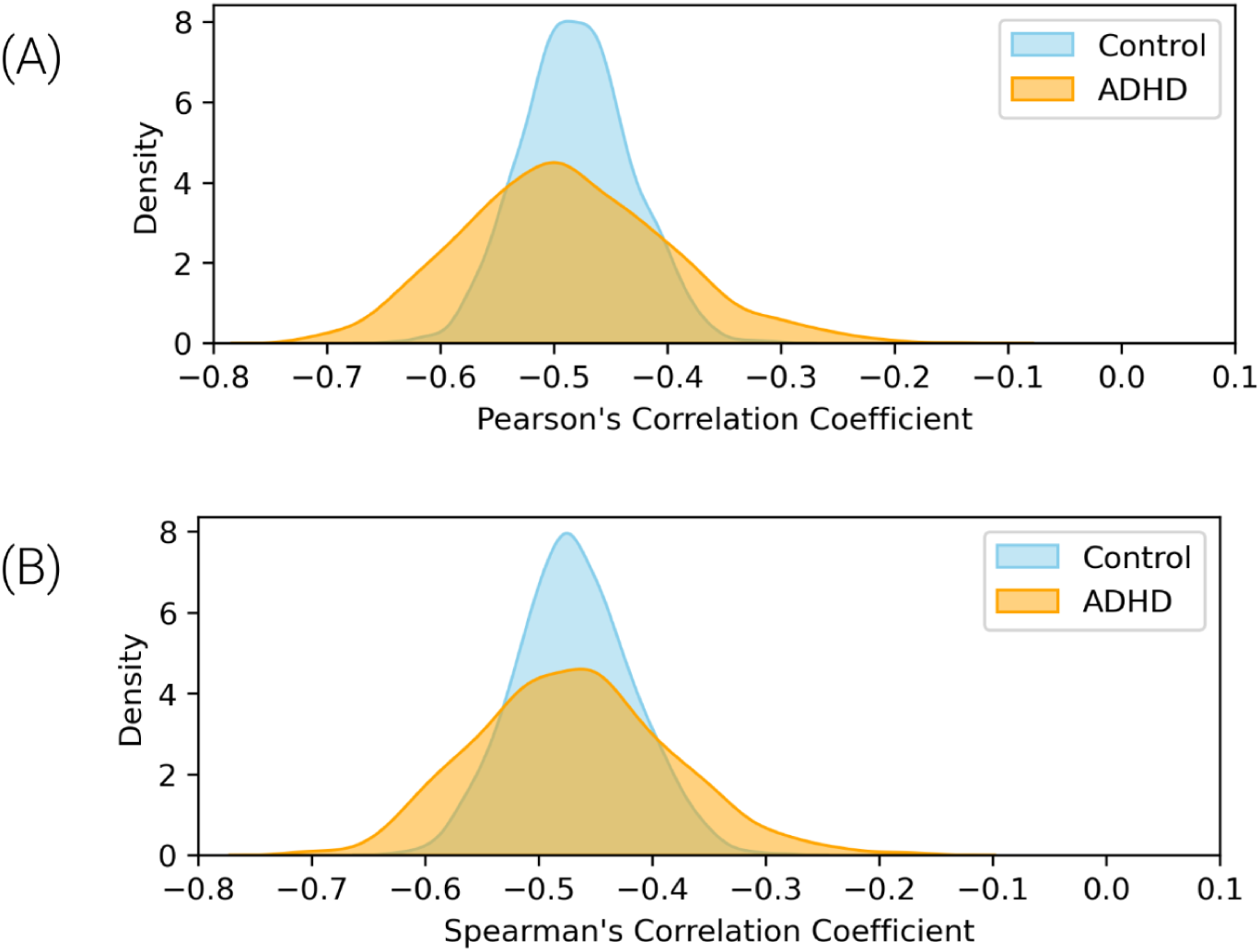
The bootstrap kernel density estimation (KDE) (*kernel = Gaussian; bandwidth = 1*), of correlation coefficients for control (blue) and ADHD (orange) groups. (A) Distribution of Pearson’s Correlation Coefficients; (B) Distribution of Spearman’s Correlation Coefficients. The overlap indicates that the presence of ADHD symptoms does not significantly affect the decrease in Ising temperature with age.

### 2.3 Confound analysis

To evaluate the influence of possible confounds, the effect of head motion was investigated on the Ising Temperate estimation since its significant impact on fMRI data (Friston et al., 1996). Thus, a linear regression was used to predict which variables could predict temperature (2 confounding variables: Max Motion (mm) and Max Rotation (degree), and Age (years) as the variable of interest), where Max Motion (mm), Max Rotation (degree) and Age (years) as independent variables and Estimated Ising temperature as the dependent variable, for each group. For the Typically Developing Children group model (R^2^= 0.19), age was the only significant variable influencing temperature, showing a negative association (β =− 0. 43, two-sided *p* < 0.001). For the ADHD symptoms group model (𝑅 = 0. 32) repeats this same pattern (β =− 0. 37, two-sided *p* < 0.001). The results are shown in Table 2. Typically Developing Children ADHD symptoms

**Table 2.** Linear regression summary (linear coefficient *β*, *p*-value and 𝑅 ) for the evaluation of head motion effects in Ising temperature estimation for Typically Developing Child and ADHD symptoms groups. A model was utilized for each group separately, using Max Motion (mm), Max Rotation (degree), and Estimated Ising temperature as features to explain the target variable Age (years).

|  | Typically Developing Children |  | ADHD symptoms |  |
| --- | --- | --- | --- | --- |
| | $\beta$ | $p$ | $\beta$ | $p$ |
| Max Motion (mm) | -0.085 | 0.24 | 0.029 | 0.73 |
| Max Rotation (degree) | -0.065 | 0.40 | -0.033 | 0.65 |
| Age | -0.434 | < 0.001 | -0.365 | < 0.001 |
| $R^2$ | 0.190 | | 0.324 | |

## 3 Discussion

This paper is based on past research on complex systems, functional connectivity, and criticality, in particular, inspired by the evidence that the brain is poised near a second-order phase transition (Fraiman et al., 2009) and that brain criticality using Ising models can also be used to investigate brain states (Ruffini et al., 2023, Kandeepan et al., 2020). The present framework assumes that the 2D Ising model can describe the empirical resting functional networks–as proposed by Fraiman et al. (2009)–while expanding this concept over neurodevelopment. Thus, this work focuses on investigating brain criticality rather than characterizing changes in the topological organization of functional networks. Critical models have been a key to understanding the emergence of collective complex neural dynamics and can be clinically relevant to many domains – including anesthesia, sleep medicine, developmental-behavioural pediatrics, and psychiatry (Jiang et al., 2018; Zimmern, 2020; Rocha et al., 2022). During the critical phase, the model achieves an equilibrium between alignment interactions and thermal excitations (entropy gain). This equilibrium gives origin to the emergence of a diverse repertoire of complex patterns. Hence, estimating the Ising temperature of rs-fMRI creates a bridge to investigate the availability of dynamical states that the system can occupy and transit in different conditions, such as neurodevelopment.

The main finding indicates a statistically significant negative correlation between age and the Ising temperature for Typically Developing Children and ADHD symptoms, suggesting that the brain gets distant from criticality to an ordered state as age increases. This result was consistent across two different orders of magnitude for the number of time points in the 2D Ising model. Previous studies support that resting-state brain dynamics stay slightly below the critical point in a sub-critical regime (Skilling et al., 2019; Priesemann et al., 2014). However, this remains controversial, with other evidence showing a slight shift to a paramagnetic regime (Ezaki et al., 2020; Ruffini et al., 2023). Different pre-processing pipelines and multiple co-factors could potentially influence the estimation, and therefore, the most valuable insights come from the overall trend. This present paper suggests a move from a disordered phase to a more ordered phase, where neural activity becomes less random and more predictable. The correlation coefficients for the groups were statistically indistinguishable from each other, suggesting that the maturation process, as measured by the Ising temperature, follows a similar trajectory in both typically developing children and those with ADHD symptoms. This could be influenced by the use of medication in ADHD subjects. For both groups, possible motion confounds were excluded using linear regression analysis.

One of the most common frameworks to infer Ising temperature is based on the PMEM (Ruffini et al., 2023; Ezaki et al., 2017) that finds the appropriate probability distribution given data constraints. These techniques also allow the estimation of the coupling matrix **J,** which enables different strengths for interactions between spins and the external magnetic field ***h***. Moreover, other studies also use the structural geometry of the brain as measured by diffusion tensor imaging to biologically inform the Ising model (Marinazzo et al., 2013; Kandeepan et al., 2020; Nuzzi et al., 2020). Even though most of these frameworks are more complex and robust, the model’s fit is constrained by the ROI number and time point length, also requiring much computational power to infer parameters for a large group of subjects with a vast number of ROIs (Ezaki et al., 2017). The present framework, based on machine learning, represents a simpler Ising model to compensate for these issues with the cost of several limitations: The coupling matrix **J** is constant for every spin, only direct neighbor interactions, the absence of the external magnetic field *h* and assume that the underlying model is capable of describing the empirical data. Despite the restrictions, the current approach can be improved and has significant differences and some advantages; it estimates the control parameter using directly from the connectivity information and has a very small computational cost after the training phase when inferring the Ising temperature from the connectivity matrix of a large group of subjects, respectively. Therefore, despite the present limitations and assumptions, the great advantage of this framework is the scalability of estimating the Ising temperature for large datasets with a large number of ROIs.

Furthermore, it also contributes to a new methodology for functional connectivity, using graph-structured simulation data to train a neural network that is used in empirical data. This methodology proposes integrating theoretical models with empirical data once both brain networks and Ising networks share the graph structure. Thus, this framework benefits from the possibility of expansion for biology-inspired Ising models (Marinazzo et al., 2013; Kandeepan et al., 2020; Nuzzi et al., 2020) and other criticality models, such as the Kuramoto model, which addresses how individual components (oscillators or neurons) interact to produce collective states (Kitzbichler et al., 2009; Ponce-Alvarez et al., 2015). This allows the GNN to learn complex data representations and extract information from the network topology to infer parameters. However, it is fundamental to address whether the simulation in which the trained machine learning method can describe the empirical data to ensure a meaningful inference.

A significant limitation of this study that must be addressed is the absence of the topological characterization of the functional connectivity networks. While the 2D Ising model effectively captures functional resting state networks through the critical temperature without incorporating topological features (Fraiman et al., 2009), the current approach may overlook essential aspects of network reconfiguration that influence the brain’s shift from disordered to more ordered states, weakening the results. Therefore, future models should aim to integrate topological metrics into the training set simulations and in post hoc analysis to ensure the completeness of the analysis.

## 4 Methods

### 4.1 Data description

For our study, we utilized the data provided by the Neuron Bureau for the Attention-deficit/hyperactivity disorder (ADHD) 200 competition. This dataset is public and was compiled from eight different centres. Resting-state fMRI data were acquired using 1.5 T scanners (detailed in Bellec et al., 2017). The dataset comprises 776 training subjects and 197 test subjects. The participants are categorized into four groups: healthy control, ADHD combined, ADHD hyperactive-impulsive, and ADHD inattentive. For the purpose of our analysis, we combined all ADHD types into a single category to focus on the binary classification between ADHD and healthy control participants. In addition to the basic fMRI data, various phenotypic information is provided for each subject, including age, gender, handedness, IQ, and ADHD type (Bellec et al., 2017). The head motion criteria, specifically head motion <3 mm translation or <3° rotation in any direction were applied as per previous studies (Tamm et al., 2006; Ge et al., 2019). Therefore, to restrict the influence of head motion, this threshold was reduced to 1.5 mm and 1.5°; thus, the final number of subjects was 242 typically developing children, and 111 subjects had ADHD symptoms. The dataset was pre-processed using the Athena pipeline for resting-state fMRI and voxel-based morphometry preprocessing (grey matter) with AFNI and FSL (Bellec et al., 2017). The data, which include time course values of BOLD signals, were obtained from the Connectome website (www.preprocessed-connectomes-project.org/adhd200/). Preprocessing steps included removing the first four time points to allow for magnetization to reach equilibrium, slice time correction, motion correction, registration at 4x4x4 voxel resolution using Montreal Neurological Institute (MNI) space, band-pass filtering (0.009 Hz < f < 0.08 Hz), and smoothing with a 6 mm FWHM Gaussian filter (for more details see Bellec et al., 2017). Each data collection site used its own scanner(s) and its own MR scanning parameters. Full details are available at ADHD-200-Webpage (<u>2011</u>). Only the first scan was considered for participants with multiple scans and selected 140 time points for every subject.

In order to minimize the risk of co-factors and co-foundings over age, the demographic information was evaluated. To remove the possibility of sex impact on age, the distribution was investigated (Male, 12.7 **±** (3.8) years, Female, 12.4 **±** (3.5) years) and also the ADHD impact on age (Typically Developing Children, 12.8 **±** (3.7); ADHD-Combined, 11.6 **±** (3.4); ADHD-Hyperactive/Impulsive, 14.4 **±** (4.9); ADHD-Inattentive 11.9 **±** (3.0) ). Furthermore, motion was also evaluated, showing the max rotation (degree) has statistically significant Pearson’s correlation value with age (Max rotation (degree), *r = -0.13*, *p = 0.01,* and Max motion (mm), *r = 0, p > 0.05*). Both distributions can be seen in Fig. 4.

**Figure 4.**
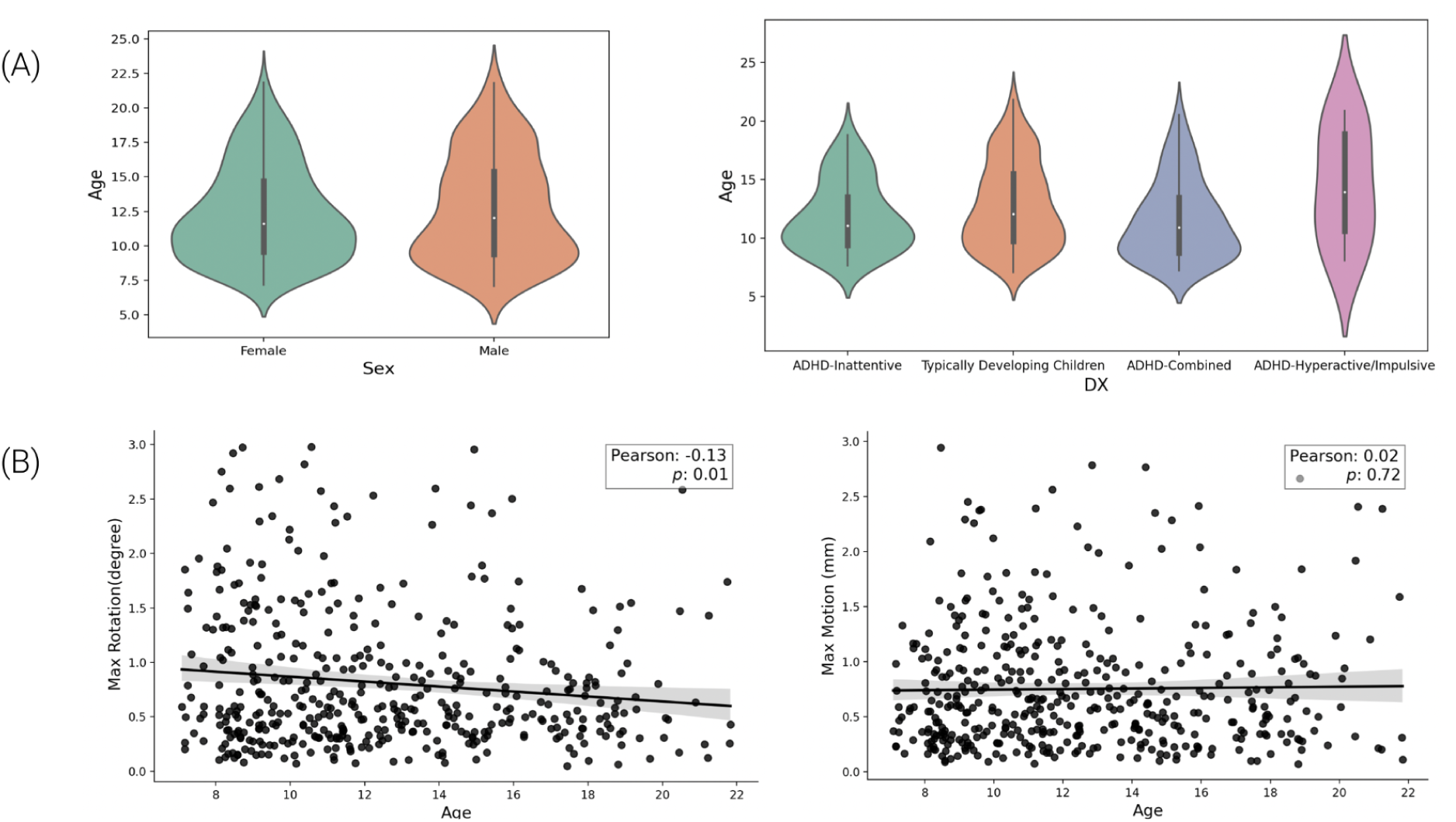
The main demographic aspects of the dataset. (A) The distribution of age across sex and the four categories of participants (Typically Developing Children, ADHD-Combined, ADHD-Hyperactive/Impulsive, and ADHD-Inattentive). (B) Scatter plot with Pearson’s correlation and p-value for Max Rotation (degree) and Max Movement (mm), the solid black line in each panel represents the linear trend between head motion and age, with a shaded region indicating the standard deviation.

To better understand the influence of the presence of ADHD symptoms in motion, the dataset was separated into two groups: Typically Developing Children and ADHD symptoms. Thus, the max motion (mm) and age for the ADHD group were the only statistically significant positive correlations (Max motion (mm), Pearson’s r = 0.27, p-value = 0.001; and Spearman’s r = 0.25, p-value = 0.001). Both distributions can be seen in Fig. 5.

**Figure 5.**
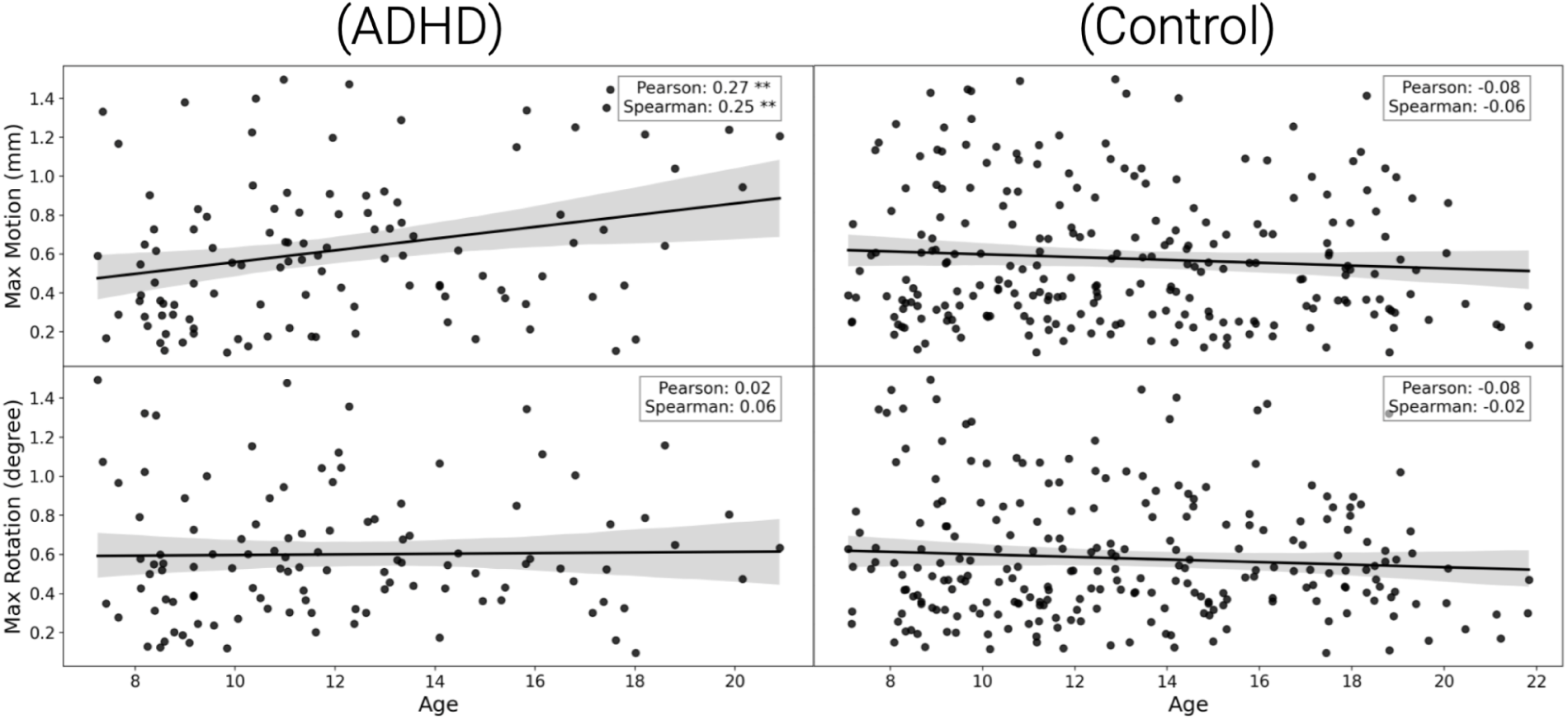
Scatter plots showing Pearson’s correlation and p-values for Max Movement (mm) and Max Rotation (degrees) as a function of age in children diagnosed with ADHD (left) and Typically Developing children (right). The top row represents Max Movement (mm), while the bottom row represents Max Rotation (degrees). Each scatter plot includes a linear trend line (solid black) with the shaded region indicating the standard deviation. Pearson’s correlation coefficient and Spearman’s rank correlation are displayed in each panel, with asterisks denoting the significance levels (* p < 0.05, ** p < 0.01, *** p < 0.001 ).

### 4.2 Image processing

The functional data was parcellated into 333 cortical regions of interest (ROIs), as defined by (Gordon et al., 2016). This parcellation was motivated by the intense changes the cortex undergoes during neurodevelopment (Baum et al., 2020) and by the intrinsic critical nature of cortical networks (Páscoa dos Santos et al., 2023). Then, we obtained the average BOLD signal of the voxels in each ROI. Pearson’s pairwise correlation of different ROIs was calculated to create the adjacency matrix A (connectivity matrix). Therefore, a brain graph is defined as **G = (V, E, A)**, with **V = {1, …, N}** a set of nodes (or vertices), E representing a set of corresponding edges, and **A** ∈ **R^NxN^** denoting the weighted adjacency matrix. One entry ***w_ij_*** of the adjacency matrix **A** would indicate the connection strength (Pearson’s Correlation) between node ***i*** and node ***j*** of graph **G**.

### 4.3 2D Ising model Simulation

The 2D Ising model describes ferromagnetic materials and demonstrates magnetization resulting from a second-order phase transition. The model consists of a grid of spins, which are described by a discrete variable that takes two values σ*_i_*= −1, 1, that is, spin “up” and spin ”down,” and the Hamiltonian gives the interaction of these spins without an external magnetic field (McCoy et al., 1973),

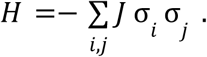

Where 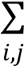 represents the sum of neighbouring spins, and J is the coupling constant that describes how strong the interaction between the spins is. Using 𝐻 = 𝐸, it is possible to obtain energy information for any state of the grid.

Given the discrete and stochastic nature of the Ising model, Monte Carlo algorithms are convenient for calculating some model estimates. More specifically, the algorithm Metropolis-Hastings (Gould and Tobochnik, 1996; Metropolis et al., 1953). The algorithm is built based on the concept of Markov chains (see Supplementary Material for more details). All simulations were performed on a lattice 𝐿 = 250, starting with all spins aligned or else randomly distributed, with Monte Carlo time steps Δ𝑡 = 𝐿 × 𝐿 to cover all sites in the network giving a chance for every spin to change its state at least once (Fraiman et al., 2009). The 2D Ising model undergoes a second-order phase transition when passing through the critical temperature 𝑇_𝑐_ ≈ 2. 269𝐽 (McCoy et al., 1973). The Boltzmann constant and the coupling constant are set to 𝑘_β_ = 𝐽 = 1.

The 2D Ising model was simulated using a lattice of 250 × 250 spins fluctuating over 200 time steps after thermal equilibrium, where each time step allows all the spins of the lattice to change their state. Once the model is indeed in thermal equilibrium, the lattice is averaged over blocks of 13 × 13 spins, in order to get a continuous time series representative of the fMRI signal (Fig. 6). Since 250 is not divisible by 13, a floor division is applied, which means extracting just the integer part of a division. This reduces the system to a smaller grid where each block is averaged, resulting in a lattice of 250//13 x 250//13 = 19 x 19, which contains 361 ROIs. Now, we calculate the correlation matrix and then remove the excess rows and columns to get 333 sizes. The full correlation matrix is initially larger (361 x 361), but the target is a smaller matrix (333 x 333). By calculating how many rows and columns to remove from each side, the code extracts the central portion of the matrix, removing 14 rows and columns from both the top and bottom, as well as the left and right, ensuring the final matrix has the desired dimensions. This helps to eliminate boundary effects. The 2D Ising model graphs were calculated by resulting in 333 nodes (the simulation and processing are open-source, available on https://github.com/Rodrigo-Motta/BRAIN_ISING_GNN).

**Figure 6.**
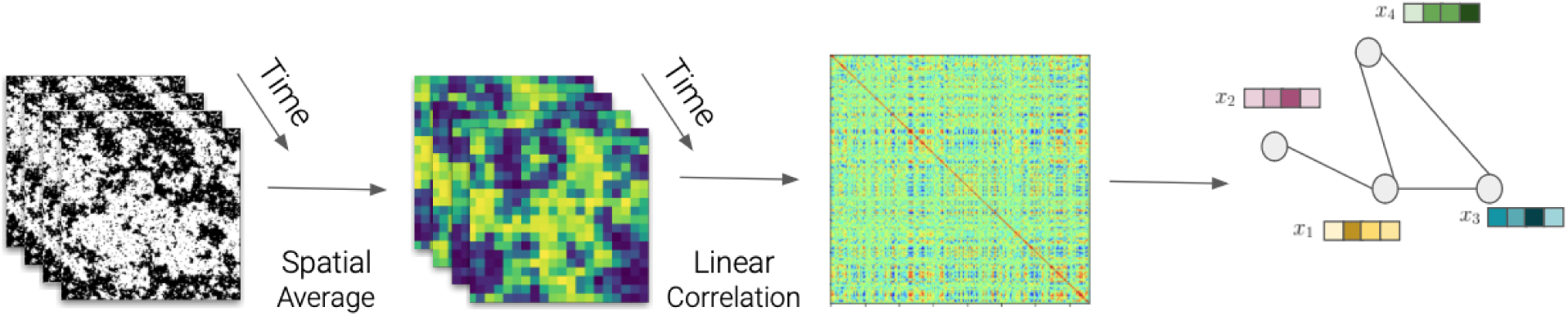
The creation of 2D Ising model networks process. Firstly, the spatial average of near spins results in a continuous version of the simulated dynamical 2D Ising Model. Then, a pairwise Pearson’s correlation is calculated over all lattices to create the graph adjacency matrix with the same dimensionality as the rs-fMRI connectivity matrix.

We simulated time points in the 2D Ising model for 200 and 2000 Δ𝑡 , finding that the Graph Neural Network performed slightly better at predicting the Ising temperature with 200 time points. Therefore, we chose to proceed with 200 time points for further analysis. Nonetheless, despite the performance differences, the results were consistent even with 2000 time points, as detailed in the supplementary materials.

To check that the 2D Ising model simulation routine was correctly employed and thermalized, in the absence of an external magnetic field, the order parameter (mean magnetization per site) was evaluated in a wide range of [1. 6, 3. 3] 𝑘_β_𝑇/𝐽. For model robustness, the evaluation was conducted for three different lattice sizes compared with the analytical solution. The chosen lattice size was selected based on less variability during the critical regime and dimensionality proximity with Gordon’s parcellation of 333 ROIs.

### 4.4 Graph Convolutional Neural Networks (GCNs)

The GCN mechanism can be described by convolution operations, non-linear activation functions, pooling, and backpropagation. Firstly, the graph convolution operation is the primary operation of a GCN, extracting topological structures from the graph. This operation is based on a message-passing operation (for more details, see Supplementary Material). These intrinsic geometric operations from GCNs allow the complex representation learning and prediction of global parameters from networks. In this work, this model learns to estimate a global property of the graph by extracting and transforming connectivity features from nodes by graph convolutional layers (Kipf & Welling, 2016).

Even though GNN operates in graph structures, it’s necessary to process the graphs for the GNN. Thus, the k-nearest neighbors (KNN) algorithm was used to determine the graph edges to connect each node and its neighbors. This approach effectively avoided problems with fitting noise and vanishing gradients, where each node represented an ROI and the node features were the connectivity with the other ROIs (Qin et al., 2022; Lei et al., 2022). A GCN was trained to calculate the Ising temperature for the brain networks, with three graph convolutional layers, a global average pooling process, and two linear layers. The activation function was the Leaky-Rectified Linear Unit (Leaky-ReLu) in order to allow negative correlation values, the dropout technique was also used, and the loss function used was the Mean Absolute Error (MAE) with L2 Regularization to avoid overfitting. The architecture can be seen in Fig. 7.

**Figure 7.**
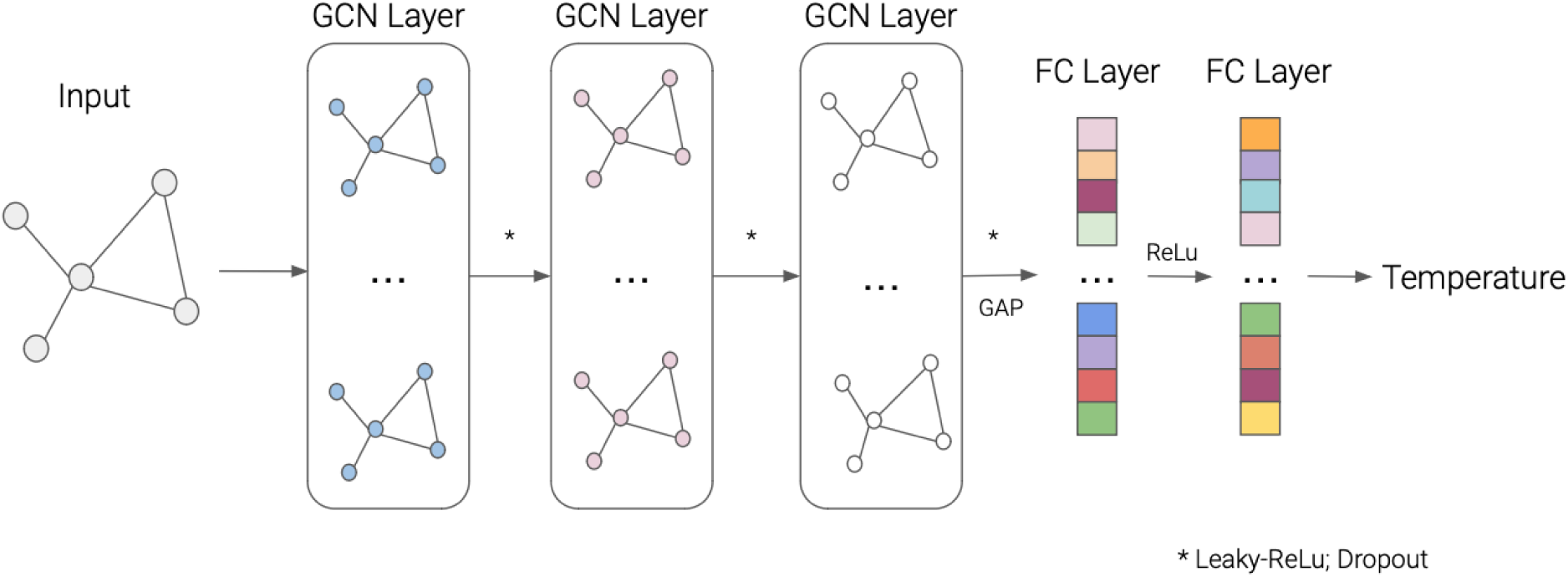
Overview of the GCN regressor architecture. Firstly, the input, a graph extracted from the ising model or empirical data, passes through 3 successive GCN layers, followed by a GAP and two successive FC layers to map the Ising temperature. Abbreviations: GAP, global average pooling; Leaky-ReLu, Leaky-Retified Linear Unit; FC, Fully Connected.

The dataset used for modeling was composed of 1500 simulations of the 2D Ising model uniformly distributed around the critical point (Temperature interval = [1. 8, 2. 5]), where 1000 simulations were used to train the network, 250 to evaluate and 250 to test. The GNN was trained over 100 epochs, using a Cycle Optimization technique; this method varies the learning rate according to triangle waves with selected maximum and minimum. In this specific work, a threshold of loss was used in the validation set to select a suitable candidate of minima to fine-tune the model in order to increase accuracy.

### 4.5 Using GNN to estimate Ising temperature

The input to a GNN is inherently a graph structure, which allows for the practical analysis of data with complex relationships. Given that both the brain and Ising model graphs share the same dimensionality, the trained GNN can seamlessly process both types of data. This dimensional consistency ensures that the model, once trained on the Ising model graphs, can be effectively used to estimate the temperature of brain graphs. This capability allows the GNN to generalize its learned representations from simulated data to real-world fMRI data. Thus, this methodology enables estimation between theoretical model parameters and empirical data. The overview of the methodology can be seen in Fig. 8.

**Figure 8.**
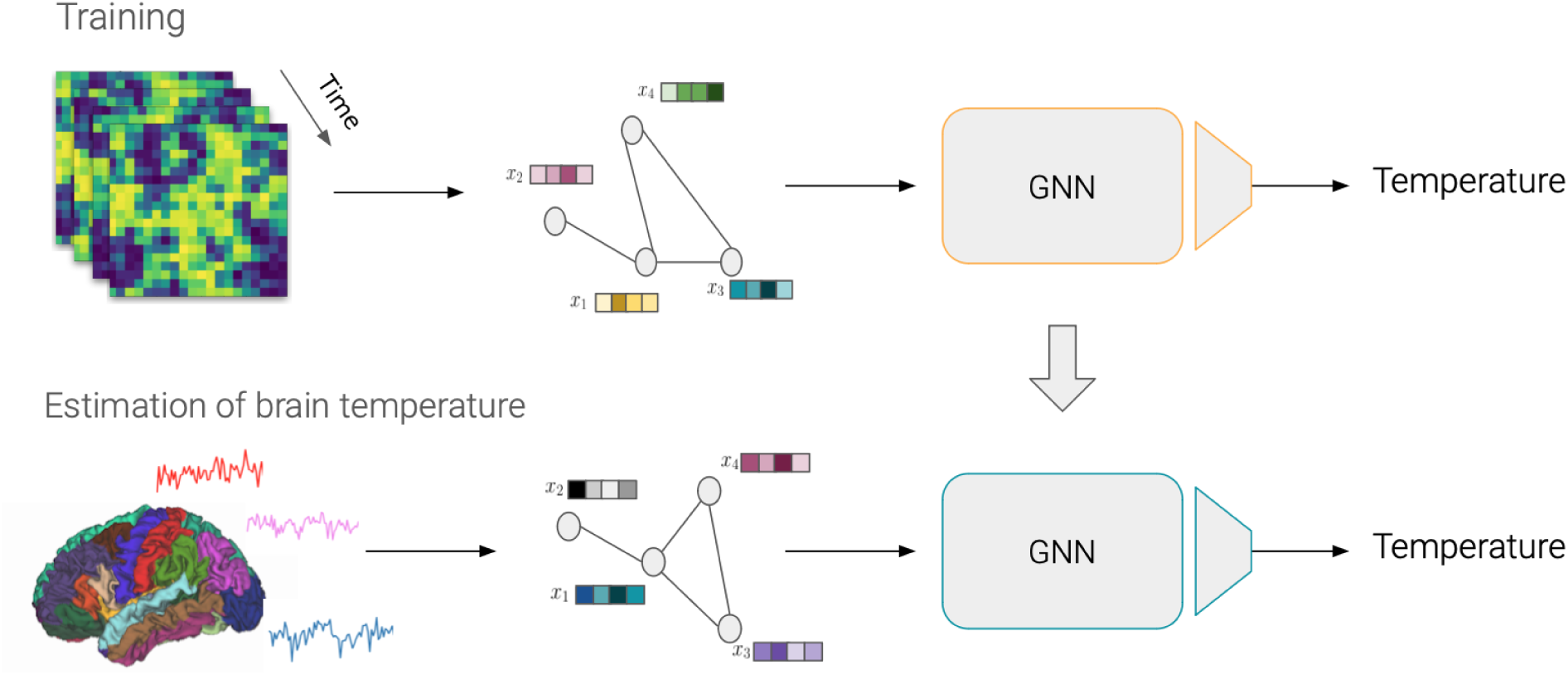
Overview of the methodology. On the top, the training phase of the Graph Neural Network (GNN) used multiple simulated Ising model networks at different temperatures to create a regressor. On the bottom, the trained GNN is used to estimate the Ising temperature using whole brain networks.

After training the Graph Neural Network (GNN) on 2D Ising model simulations near criticality ([1.8, 2.4] k_B_T/J), hyperparameter tuning was performed on the validation set to optimize its performance. The GNN demonstrated high efficiency in predicting temperature on the test set, achieving a mean absolute error 𝑀𝐴𝐸 = 0. 04 and an 𝑅 = 0. 92.

### 4.6 Statistical Analysis

Firstly, to evaluate the overall trend of the temperature over age, Pearson’s correlation and Spearm’s rank correlation were used. A bootstrap was done to check the statistical differences between correlation coefficients between groups (ADHD and Control). The bootstrap analysis was implemented using 1000 resamples, following the bias-corrected and accelerated bootstrap (see Details in Efron et al., 1996). In each repetition, the correlation coefficient was computed, and the resampling was performed with replacement, where the average percentage of subjects excluded in each resample was 36.8%, following the probability of not being chosen. The confidence level was set at 95%, meaning the resulting bootstrap confidence interval covers the actual correlation coefficient approximately 95% of the time.

To check possible differences in temperature between different age groups, the data was separated into seven groups of 2 years (7-9, 9-11, 11-13, 13-15, 15-17, 17-19, 19-21). The percentage distribution of data across the age groups is as follows: The age group 9-11 has the highest percentage at 20.17%, followed closely by the 11-13 group at 19.75%. The 13-15 group accounts for 16.39%, while the 7-9 group represents 15.55%. The age group 17-19 makes up 15.13%, and the 15-17 group comprises 9.66%. Lastly, the 19-21 age group has the smallest percentage, accounting for 3.36% of the total data.

## Data and Code Availability

All software and procedures concerning the analysis were detailed in the methods, and supplementary materials and scripts are available on GitHub: https://github.com/Rodrigo-Motta/BRAIN_ISING_GNN. The dataset, which includes time course values of BOLD signals, was obtained freely on (http://fcon_1000.projects.nitrc.org/indi/adhd200/).

## Author Contributions

Conceptualization: R.C., J.R. Data curation: R.C., J.R. Methodology: R.C., W.P., J.R. Investigation: R.C., W.P., J.R. Resources: R.C., J.R Formal analysis: R.C., J.R Visualization: R.C. Writing—original draft: R.C., J.R. Writing—review & editing: R.C., J.R. Project administration: J.R. Software: R.C. Visualization: R.C. Supervision: W.P., J.R. Funding acquisition: R.C., J.R.

## Acknowledgements

This study is supported by the São Paulo Research Foundation (FAPESP) Grants 2021/05332-8, 2023/02616-0, 2023/02538-0, 2024/00861-0, and 2024/09675-5.

