## Supplementary Material for "A Graph Neural Network Approach to Investigate Brain Critical States Over Neurodevelopment"

### Impact of the number of fluctuations

To evaluate whether the number of fluctuations affects the results, we examined the outcomes for T=200 and T=2000. In the 2D Ising model, the number of fluctuations affects the measurement of thermodynamic properties, specifically, with T=200 capturing fewer fluctuations, representing shorter system dynamics, while T=2000 reflects a more extensive sampling of the system's behavior over time. Despite this difference, we observed a negative correlation between predicted temperature and age for both control and ADHD groups. Interestingly, the Graph Neural Network performs similarly on the simulation test set for both time points, with slightly better performance for T=200 (T=200: MAE=0.04, $R^2$=0.92; T=2000: MAE=0.06, $R^2$=0.86).

| | | *Pearson's Correlation* | *p-value* | *Spearman's Correlation* | *p-value* |
|---|---|---|---|---|---|
| *Control* | *T=200* | *-0.48* | *$p < 0.0001$* | *-0.47* | *$p < 0.0001$* |
| *ADHD* | | *-0.49* | *$p < 0.0001$* | *-0.47* | *$p < 0.0001$* |
| *Control* | *T=2000* | *-0.25* | *$p < 0.001$* | *-0.19* | *$p < 0.01$* |
| *ADHD* | | *-0.20* | *$p < 0.01$* | *-0.19* | *$p < 0.01$* |

**Table A.** The correlation coefficients and p-values for both Pearson's and Spearman's correlations between predicted temperature and age, separately for Control and ADHD groups, under two different time points (T=200 and T=2000). Negative correlation coefficients indicate an inverse relationship. Results are statistically significant at varying levels, with stronger correlations and lower p-values observed at T=200 compared to T=2000. Pearson's correlation quantifies linear relationships, while Spearman's correlation captures monotonic relationships.

### Temperature and global graph metrics

Despite the difficulty of describing changes in network topology in Ising Temperature, relationships can be explored by verifying Pearson's correlation with other global graph metrics within the groups. Thus, some main graph topological properties were used: Average clustering, average betweenness centrality, average path length, average degree, and average degree standard deviation. Therefore, the temperature change strongly correlates to all measures except for average betweenness centrality in both groups.

There are distinct differences between empirical data (Control and ADHD) and Ising simulations in how functional connectivity responds to temperature changes. While the Ising model shows consistent trends—such as increased clustering and path length with temperature changes—the empirical data exhibit weaker or opposing relationships, particularly in ADHD networks, which show reduced correlations overall. This discrepancy highlights the Ising model's limitations in fully capturing real-world functional connectivity dynamics. The simulations' more significant, more systematic changes suggest that the Ising model may better reflect idealized, theoretical network behaviors, whereas additional biological and structural complexities influence empirical networks. Moreover, the significance of the average degree standard deviation decreases with increasing Ising temperature on empirical data and simulations, reflecting a homogenization of node connectivity in the network. This aligns with the theoretical expectation that an increase in temperature drives the system toward a more disordered, less differentiated state, where node connections become more uniform.

| | *Control* | | *ADHD* | | *Ising* | |
|---|---|---|---|---|---|---|
| *Metric* | *r* | *p* | *r* | *p* | *r* | *p* |
| *Average Clustering* | *-0.34* | *<0.0001* | *-0.20* | *<0.05* | *0.41* | *<0.0001* |
| *Average Betweenness Centrality* | *-0.04* | *0.2* | *-0.05* | *0.4* | *-0.16* | *0.1* |
| *Average Path Length* | *0.09* | *0.4* | *-0.07* | *0.4* | *0.66* | *<0.0001* |
| *Average Degree* | *-0.35* | *<0.0001* | *-0.23* | *<0.05* | *0.43* | *<0.0001* |
| *Average Degree Standard Deviation* | *-0.54* | *<0.0001* | *-0.35* | *<0.0001* | *-0.73* | *<0.0001* |

**Table B.** Pearson's correlation between Ising Temperature changes and global graph metrics for the empirical data (Control and ADHD) and Ising simulations. The table presents the metrics, Pearson correlation coefficients ( $r$ ) and the corresponding p-values ( $p$ ) for the relationship between changes in the estimated Ising Temperature and various graph topological properties, including Average Clustering, Average Betweenness Centrality, Average Path Length, Average Degree, and Average Degree Standard Deviation.

### 2D Ising model simulation

The 2D Ising model describes ferromagnetic materials in a simplified way using just one free parameter (Temperature). This model shows magnetization resulting from a phase transition, and it consists of a grid of spins, which are described by a discrete variable that takes two values $\sigma_i = -1, 1$, and the interaction of these spins given by Hamiltonian in the absence of external magnetic field (McCoy et al., 1973),

$$H = -\sum_{i,j} J \sigma_i \sigma_j$$

J is the coupling constant and $\sum_{i,j}$ is the sum of the nearest neighboring spins. When set $H = E$, it is possible to obtain energy information for any lattice configuration or state of the grid. To solve the 2D Ising model, the algorithm Metropolis-Hastings is used considering an isolated system (Gould and Tobochnik, 1996; Metropolis et al., 1953). The algorithm is built based on the concept of Markov chains, which is a stochastic model that describes a sequence of possible events in which the probability of each event depends only on the state reached in the previous event. A Markov process is solely based on the transition probabilities $P(X_{n+1} = \mu \mid X_n = \nu)$, which are the probability of going μ from ν. In this way, the entire process can be reduced to a matrix ***P*** that contains the information to go from one state to another in one step. From these properties, it follows that for a matrix ***P***, the Markov process will have a distribution stationary station that follows the balance equation,

$$\frac{P(\mu|\nu)}{P(\nu|\mu)} = \frac{\pi(\mu)}{\pi(\nu)}.$$

That is, the probability of transition from μ ν being in ν is equal to the probability of transition from ν μ being in μ. Rewriting this relation as

$$P(\mu|\nu)\pi(\nu) = P(\nu|\mu)\pi(\mu).$$

To simulate the Ising model, it is enough to know $\pi(x)$. Using the canonical and immediate ensemble,

$$\pi(x) = p(x) = \frac{1}{Z}e^{-\beta E_x}$$

where $Z = \Sigma\, i\, exp(-\beta Ei)$ is the partition function and $\beta = 1/(k_\beta T)$ (McCoy et al., 1973). This way, the balance equation is achieved,

$$\frac{P(\mu|\nu)}{P(\nu|\mu)} = \frac{\pi(\mu)}{\pi(\nu)} = e^{-\beta\Delta E}$$

where $\Delta E = (E_\mu - E_\nu)$. Changing the value of a single variable changes its state, or the $\Delta E$ flip of a spin in position $(i, j)$ will be just the energy difference in its neighbors since it will not change the energy at any other point on the grid. Therefore, considering the power before the flip

$$E_n = -J\sigma_{(i,j)}[\sigma_{(i+1,j)} + \sigma_{(i-1,j)} + \sigma_{(i,j+1)} + \sigma_{(i,j-1)}].$$

After rotating the spin $\bar{\sigma}_{(i,j)}$ the energy will be given by

$$E_{n+1} = -J\bar{\sigma}_{(i,j)}[\sigma_{(i+1,j)} + \sigma_{(i-1,j)} + \sigma_{(i,j+1)} + \sigma_{(i,j-1)}].$$

Therefore, the variation in energy of rotating the spin

$$E_{flip} =- 2J\sigma_{(i,j)}[\sigma_{(i+1,j)} + \sigma_{(i-1,j)} + \sigma_{(i,j+1)} + \sigma_{(i,j-1)}].$$

With the balance equation $E_{flip}$ calculated, it is possible to create an iterative routine to simulate the model.

(1) First, we call a state, then we rotate a spin randomly on the grid and calculate the energy $E_{flip}$;

(2) If the energy increases, we define $P(\mu \rightarrow \nu) = 1$, by the equilibrium equation, the spin inverts again mind if $P(\mu \rightarrow \nu) = exp(- \beta\Delta E)$;

(3) If the energy decreases, then $P(\nu \rightarrow \mu) = 1$ the state remains with the spin turned;

(4) Repeat this process for all spins.

The simulations were performed on a lattice $L = 250$, starting with all spins aligned or else randomly distributed, with Monte Carlo time steps $\Delta t = L \times L$ to cover all sites in the network, giving, on average, the chance of each spin of the lattice changing its state (Fraiman et al., 2009). Furthermore, as the grid is finite, the spins that are located on the edges are linked by the periodic boundary condition, in which the spins on the right edge interact with the left edge spins. With Onsager's analytical solution for the case $h = 0$ with no external magnetic field, it was discovered that the 2D Ising model undergoes a second-order phase transition when passing through the critical temperature $T_c = 2J/( ln(1) + \sqrt{2}) \approx 2.269J$ (McCoy et al., 1973). The Boltzmann constant and the coupling constant are set to $k_\beta = J = 1$.

In the simulation, a (250 x 250) lattice is divided into blocks of size 13. Since 250 is not divisible by 13, a floor division is applied, which means extracting just the integer part of a division. This reduces the system to a smaller grid where each block is averaged, resulting in a lattice of 250//13 x 250//13 = 19 x 19, which contains 361 ROIs. Now, we calculate the correlation matrix and then remove the excess rows and columns to get 333 ROIs. The full correlation matrix is initially larger (361 x 361), but the target is a smaller matrix (333 x 333). By calculating how many rows and columns to remove from each side, the code extracts the central portion of the matrix, removing 14 rows and columns from both the top and bottom, as well as the left and right, ensuring the final matrix has the desired dimensions. This helps to eliminate boundary effects.

### Graph Neural Networks (GNNs)

In order to define a graph convolution operation, it's essential to define a graph as $G = (V, E, A)$, where $V$ denotes a set of $|V| = N$ nodes (or vertices), $E$ represents a set of corresponding edges, and $A \in R^{NxN}$ denotes the *weighted adjacency matrix*. One entry $\omega_{nm}$ of the adjacency matrix $A$ would indicate the Pearson's Correlation between the time series of the node $n$ *a*nd node $m$ of a graph $G$. The graph must also have a feature matrix $X$ in order to be processed by a GCN, which, in this specific case $X$, is also the connectivity matrix; therefore, this graph can be fully explained by the adjacency matrix.

The GCN mechanism can be described by convolution operations, non-linear activation functions, pooling, and backpropagation. Firstly, the graph convolution operation is the primary operation of a GCN, extracting topological structures from the graph. This operation is based on a message passing operation, which can be simply defined as

$$f(X) = \sigma(\overline{A}XW)$$

Where $\overline{A}$ is the normalized adjacency matrix**,** $X$ the feature matrix, $W$ **is** the learnable parameter matrix, and σ is a non-linear function (Defferard et al., 2016; Kipf & Welling, 2016). The normalized adjacency matrix solves the problem of feature values for nodes with many neighbors; this summation embedded in the matrix multiplication can overestimate nodes with a high degree. Therefore, it's common to solve this problem using the degree matrix using a symmetric normalization as follows:

$$\overline{A} = D^{(-1/2)} A D^{(-1/2)}$$

Where $D$ is the diagonal degree matrix of $A$. This normalized adjacency matrix $\overline{A}$ is then multiplied by the node feature matrix $X$ and the learnable matrix $W$, propagating node features across the graph and updating $X$ for $f(X)$. Each layer updates the node features with dimensionality reduction in the bottleneck design. The convolutional operation in graphs is similar to the convolution operation in CNNs but is applied to the graph's adjacency matrix instead of a regular grid. Unlike filters in Convolutional Neural Networks (CNNs), our weight matrix $W$ is unique and shared among every node.

Equation of state calculations by fast computing machines. *Journal of Chemical Physics, 21*(6), 1087. https://doi.org/10.1063/1.1699114
